# Validation of DeepLabCut as a tool for markerless 3D pose estimation

**DOI:** 10.1101/2022.03.29.486170

**Authors:** Veronika Kosourikhina, Diarmuid Kavanagh, Michael J. Richardson, David M. Kaplan

## Abstract

Deep learning-based approaches to markerless 3D pose estimation have taken neuroscience by storm. Yet many of these tools remain unvalidated. Here, we report on the validation of one increasingly popular tool (DeepLabCut) against simultaneous measurements obtained from a reference measurement system (Fastrak) with well-known performance characteristics. Our results confirm close (mm range) agreement between the two, indicating that deep learning-based approaches can be used by the research community with confidence.

## Introduction

Recent advances in computer vision and machine learning have catalyzed the development of powerful tools for markerless 3D pose estimation (Mathis et al. 2020, Mathis & Mathis, 2020). Although these tools afford promising new opportunities for rapid, efficient quantitative measurement of animal and human behavior in neuroscience and a range of other fields (Datta et al. 2019), their accuracy and reliability have not been rigorously established. In this brief communication, we report on the validation of one increasingly popular open-source deep learning-based pose estimation tool – DeepLabCut (DLC; Mathis et al. 2018; Nath et al. 2019). We systematically compared 3D kinematic data generated via a DLC-based workflow using video frames collected from two cameras positioned at different viewing angles against simultaneous measurements obtained from a reference measurement system with well-known performance characteristics. We selected Fastrak (Polhemus, Vermont, USA), an electromagnetic 3D motion tracking system, as our reference system because of its high accuracy (static position accuracy: 0.76 mm RMS) and its wide use in human neuroscience.

## Results

To evaluate DLC tracking performance with naturalistic 3D human arm motion, participants performed two generic upper limb motor tasks — a standard center-out reaching task and a more freeform, zigzag movement task. Agreement between concurrent DLC and Fastrak data was determined by calculating bias and limits of agreement (LOA) using the Bland-Altman method (Bland & Altman, 1999), root mean square error (RMSE), and conducted a time series analysis.

We found extremely close agreement between DLC and Fastrak (summarized in Figure 2). Both bias estimates and RMSE indicate close agreement, with an average bias value of 0.8mm and an average RMSE of 1.4mm. The mean LOA range was 5.3mm, suggesting that most datapoints from DLC and Fastrak differ by no more than half a centimeter. Relative to Fastrak, DLC generally underestimated the distance from sensor position to reference point by 0.3-2.3mm (mean bias, Fig 2a). This was the case for all distance measures, except X-axis (left-right motion in camera view), where DLC overestimated the distance by 0.6-0.75mm. The overall pattern of bias was similar for center-out and zigzag tasks, but there were notable differences between finger and tool versions of the task. When the sensor was attached to the pointing tool, DLC showed more bias when estimating depth (Y-axis) than when the sensor was attached to the finger pad. It is not clear why DLC was more accurate when attached to the finger pad. The ranges of limits of agreement varied from 3.6mm to 7.0mm (i.e., from ±1.8mm to ±3.5mm relative to bias). Mean limits of agreement with respect to mean bias estimates are presented in Figure 2b. Limits of agreement vary more between trials than bias estimates (Fig 2c and Fig 2a) – suggesting that it may be possible to get better agreement in principle, but not consistently so.

**Figure 1:**
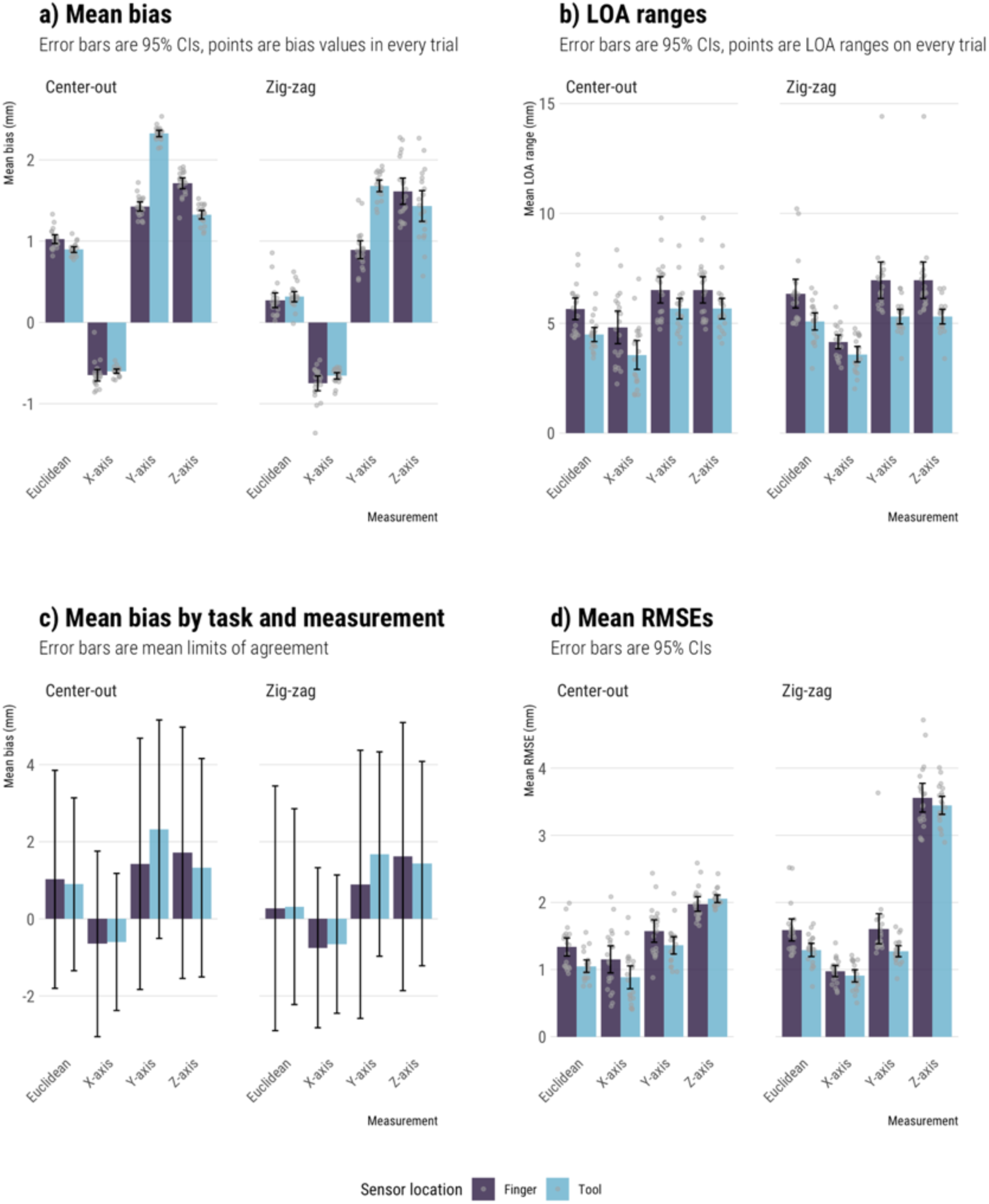
(A) Mean bias across the twenty trials, for each of the four tasks. Error bars are 95% confidence intervals and grey points are bias values for each trial (same for graphs C and D). (B) Ranges of limits of agreement across trials, for each of the four tasks. (C) Mean bias (bars) and limits of agreement (error bars) presented together. (D) Mean RMSE values across trials, for each of the four tasks.

**Figure 2:**
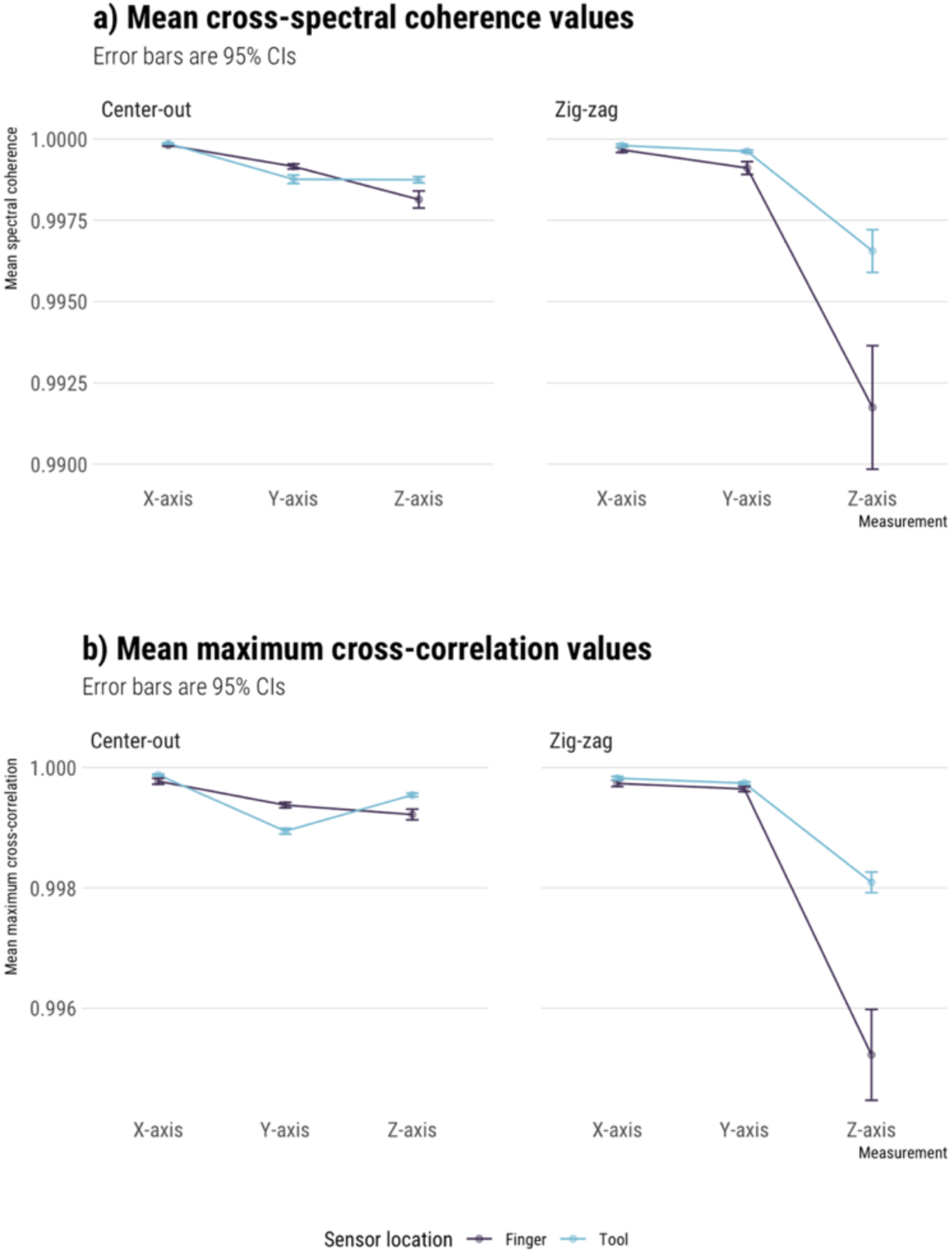
(A) Mean cross-spectral coherence values for all tasks. Error bars are 95% confidence intervals. (B) Mean cross-correlation values for all four tasks. Error bars are 95% confidence intervals.

RMSE values were similar to the bias estimates. Across the four tasks, RMSEs ranged from 0.9mm to 3.6mm, with the largest being for the Z-axis of the zigzag task (3.4mm for finger and 3.6mm for tool version of the task). In the center-out task, Z-axis estimates showed relatively worse agreement as well. Tool versions of the tasks consistently had slightly smaller RMSEs than the finger versions (except on center-out task’s Z-axis).

Both cross-spectral coherence and cross-correlation analysis revealed that there was excellent temporal agreement between the Fastrak and DLC recordings (Fig 3). Both indices have a maximum positive covariance value of 1, and the analysis resulting in values >0.99 for all three axes and across all tasks. As with other analyses, a minor reduction in temporal covariance was observed for Z-axis in the zigzag task, particularly for the finger version of the task. Overall, however, the results indicate that DLC preserves the temporal structure of human motion with the DLC and Fastrak recordings have very similar and synchronized motion patterns over time.

**Figure 3:**
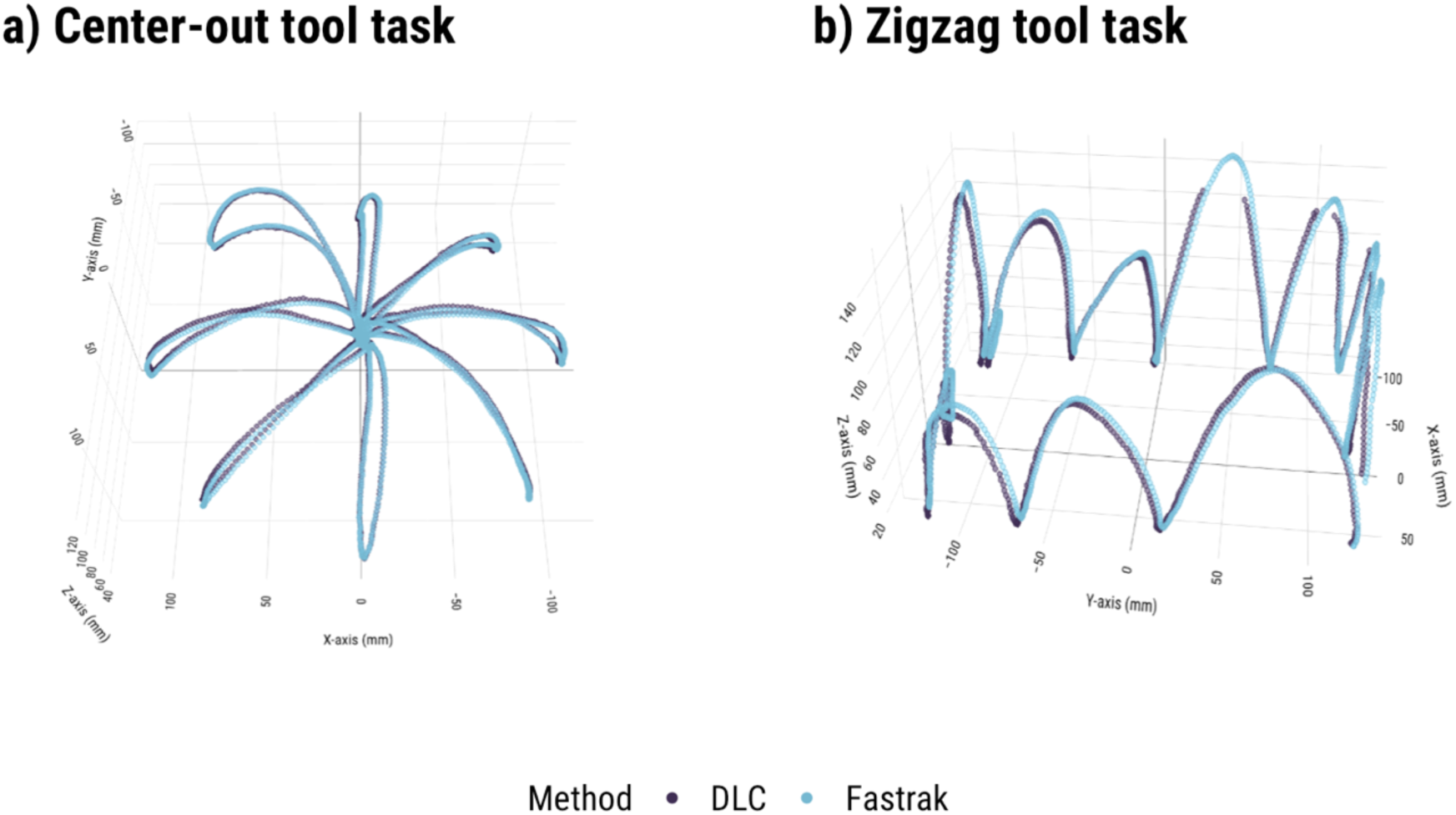
(a) Example of raw data from trial 1 of Center-Out (tool) task; (b) Example of raw data from trial 1 of Zigzag (tool) task.

## Discussion

Measurement is a fundamental aspect of modern science (Thomson 1889). When new measurement techniques or methods are adopted by a scientific field, careful validation is critical to ensure that these new methods are reliable and accurate. One commonly used approach to method validation – called the gold standard or reference system approach – involves confirming that the new method agrees within acceptable limits with an established field-standard method (Bland & Altman 1983, 1986; Choudhary & Nagaraja 2018).

Although deep learning-based approaches to markerless 3D pose estimation have taken neuroscience by storm, unfortunately, many of these tools remain unvalidated. Here, we report on the validation of one increasingly popular tool (DeepLabCut) against simultaneous measurements obtained from a reference measurement system (Fastrak) with well-known performance characteristics. We found that under tightly controlled experimental conditions, DLC can achieve extremely close (mm range) agreement with field-standard motion tracking systems such as Polhemus Fastrak, with 95% of the differences between Fastrak and DLC falling below 1 cm. The agreement was generally similar across the entire filming area, as suggested by similar patterns in agreement measures in both center-out and zigzag tasks. Despite greater bias along the Y-axis in the tool version of both tasks, bias was similar between tool and finger versions along other dimensions. Limits of agreement were smaller for all tool task data. This suggests that overall, there was a small, yet slightly greater cost in accuracy when tracking more naturalistic motion. However, this cost appears to be well within acceptable levels for most tracking and task scenarios.

Whether the degree of agreement reported in this study is acceptable will depend on the specific needs of the researcher. We found the calibration step to be the least reliable element of the DLC pipeline and getting accurate measurements compared to ground truth may require several attempts and iterative adjustment of the camera calibration settings to get the desired result. Importantly, care must also be taken to synchronize the video recordings (see supplementary methods). If done well, however, excellent 3D kinematic data can be obtained using DLC with only a minor reduction in accuracy compared to electromechanical tracking systems like Fastrak. For many applications, these slight costs in accuracy will be small compared to those imposed by having to use wired sensors or contend with noisy data due to the presence of metal near, in, and around the workspace.

## Materials and Methods

### Experimental tasks

The task set for the validation consisted of two generic, representative upper limb motor tasks — a standard center-out reaching task and a more freeform, zigzag movement task, which was specifically chosen to evaluate DLC tracking performance with naturalistic 3D human arm motion. To ensure the validation results were not effector specific, we ran two versions of the task set – one performed with the index finger and another with a pointing tool (two thin conjoined wooden dowels). For each task, all trials were completed by one researcher. In the center-out task, the researcher made reaches from a central starting position to one of 8 targets arrayed around a circle (∼12cm radius) with 45° spacing (Fig 3a). In the zigzag task, the researcher made movements around the visible edge of the testing area in a vertically oriented zigzag motion (Fig 3b). 20 trials were performed for each version of the 2 tasks (20 seconds of data recorded per trial in the center-out task and 10 seconds per trial in the zigzag task).

### Experimental setup

Data was collected with Polhemus Fastrak (Micro Sensor 1.8) and two cameras (Blackfly S BFS-U3-04S2M). The filming area was marked with a Lego mat attached to a table. A Fastrak transmitter was attached to the underside of the table, such that the entire filming area stayed within one hemisphere of the transmitter (Y+). The cameras were placed on stands attached to a platform at one side of the filming table. Edges of the stands were 30cm apart and camera lenses were ∼5cm above the table surface. The center of the filming area was ∼60cm from the lenses. A laser pointer was attached on top of each camera to verify that the cameras were pointed at the same central point a few centimeters above the filming area during all recordings. Note that cameras had to be positioned with minimal pitch relative to the filming area, because the calibration process does not handle camera pitch well and produces 3D coordinates that appear to be tilted towards the cameras (i.e., Z-axis coordinates increased disproportionately further away from the cameras).

To obtain concurrent data, we used DLC to track the position of a Fastrak microsensor. In the tool version of the tasks, the sensor was attached to the ridge formed between two joined dowels. To improve the accuracy and precision of DLC labelling, crosshairs were drawn over the center of the microsensor. In the finger version of the tasks, the sensor was attached to the pad of the index finger. Attaching sensor to the finger pad allowed easier simultaneous access to both the filming area and the computer with camera controls. Both DLC and Fastrak were set up to have the same origin point (one corner of a Lego brick located in the center of the filming area). Both systems measured the position of the sensor relative to the point of origin in three dimensions (in camera view, X-axis was left-right, Y-axis was depth, and Z-axis was up-down). Euclidean distance was calculated as a summary metric.

### Frame synchronization

Obtaining accurate 3D data and accurate comparisons with Fastrak data requires the video recordings to be synchronized across the two cameras and with Fastrak. Synchronization between the two cameras and Fastrak was achieved as follows. Camera 1 was controlled directly from the computer, and Camera 2 and Fastrak were set up to start and stop recording on signal from Camera 1. Cameras 1 and 2 were connected via a GPIO synchronization cable to a digital I/O device (NI USB-6501), which was in turn connected to the computer. Fastrak was also connected to both the computer and the I/O device in order to detect the signal that Camera 1 sent to trigger Camera 2 to start and stop recording. All three devices recorded with the same frequency (120 frames per second for the cameras, 120Hz for Fastrak), allowing for maximum temporal synchronization between data points from Fastrak and DLC. In some cases, there was a small lag in stopping the Fastrak recording (∼8 data points) – only the first 1200 or 2400 points (depending on task) were analyzed. To make the data directly comparable, we set Fastrak’s reference point to match the origin point used for camera calibration (see Calibration section). Fastrak’s settings and recording were controlled with a Matlab script, and the cameras were controlled with Multi-Pyspin (https://github.com/justinblaber/multi_pyspin). Multi-Pyspin’s code was edited to collect diagnostic information and to optimize the frame acquisition process and improve synchronization.

### Camera calibration

Accurate 3D pose estimation requires precise camera calibration, which establishes the relative location and various parameters of each camera such as the focal length and lens distortion. We chose to use easyWand over some other available alternatives (3D DeepLabCut, Anipose) as we were not able to produce reliable calibrations with these tools. We found the calibration step to be crucial to ensure reliable agreement between DLC and Fastrak data. Although all tools could produce good relative position data, it was difficult to get accurate estimates of absolute position (i.e., data looked to be the right shape, but not located where we would expect it to be in the filming space).

We used the easyWand software tool (Theriault et al., 2014), to calculate direct linear transformation coefficients and convert 2D pixel coordinates into real 3D spatial coordinates. We filmed a set of calibration videos with the necessary points labelled in DLC and used the pair with the lowest reprojection error for calculating the 3D coordinates used in the current analyses.

To calculate direct linear transformation (DLT) coefficients, easyWand requires a dataset with 2D coordinates for a set of two “wand points” and four “axis points” from each camera. Wand points are two points positioned at a fixed distance from each other that are moved throughout the volume of interest. We used the same joined wooden dowels as in the tool task, using the crosshairs over the Fastrak sensor as one point, and another set of crosshairs marked 10cm away as the second point. The axis points (one origin point and three points for each of the axes) allow aligning of the data to a particular set of axes. To get precise axis points, we placed a Lego cube and an additional Lego brick in the center of the filming area, and labelled three corners of the cube and one corner of the separate brick as the axis points. All three axis points were 31mm away from the origin point.

We filmed three pairs of calibration videos (20 seconds each), making waving motions with the dowel throughout the volume of interest. After obtaining 2D coordinates for the wand and axis points with DLC, we replaced points with <.95 likelihood with NaNs and inverted Y-axis values (by subtracting raw coordinates from image height in pixels). The latter was done to change coordinates into a bottom-oriented system, to match that apparently used by easyWand. Using raw values has consistently produced nonsense results. We saved every 10th datapoint for the list of wand-point coordinates to use in the calibration (producing ∼120-150 usable points), because using the entire set (2400 points) made it more difficult to get accurate calibrations. For axis points, easyWand only takes one coordinate point for each, so we calculated median values from our full dataset to get Origin, X, Y, and Z-axis reference point coordinates.

Intrinsic camera parameters were included to improve calibration quality: image height (540px), image width (720px), focal length (1450px), and principal point (360px). For the calibration used to acquire current data, we set the calibration to use focal length only, and applying all lens distortion coefficients. We also excluded outliers until a reprojection error of .28 was achieved with sensible estimates of camera positions. Original reprojection error (.54) was not overly large, but piloting suggested that reprojection errors <.30 are needed to achieve this degree of agreement with our setup.

### Data processing

After all data was collected, we checked timestamps from both videos and Fastrak to confirm that the correct number of data points were recorded at approximately equal time intervals. No issues were observed. To train a DLC network, we labelled 20 frames from each of two videos (camera 1 and 2) from one randomly selected trial of each task (center-out tool: trial 3; zigzag tool: trial 11; center-out finger: trial 11; zigzag finger: trial 6). Frame labelling and network training was done with default DLC settings, following the published protocol (Nath et al., 2019).

To convert 2D pixel coordinates into 3D spatial coordinates, we processed our data with a Matlab script adapted from another markerless tracking software that accepts calibration coefficients in easyWand’s format (DLTdv7, Hedrick, 2008; https://github.com/tlhedrick/dltdv). After aligning 3D data with Fastrak, we excluded any points that had <.95 likelihood in the original 2D pixel format. Plotting the remaining data as 3D scatterplots showed an apparent glitch in the Fastrak data for zigzag tasks, where its trajectory clearly deviated from the real motion path (and therefore DLC data) in one small part of the filming area. This segment of the data (X ±40mm, Y < -120mm, and Z < 25mm) was excluded from further analyses.

### Statistical analyses

To evaluate agreement between concurrent DLC and Fastrak data, we calculated bias and limits of agreement (LOA) using the Bland-Altman method (Bland & Altman, 1986, 1999), root mean square error (RMSE), and conducted a time series analysis. The Bland-Altman method is a common approach to quantify the agreement between two systems of measurement (Barnhart, Haber & Lin, 2007), and RMSE has been used as a metric of agreement in several similar motion-tracking measurement agreement studies (Niechwiej-Szwedo et al., 2018; Linke et al., 2018; Harsted et al., 2019). Bias and RMSE estimate the average difference in measurements between DLC and Fastrak, whereas limits of agreement specify the interval that 95% of measurement differences fall within. For the time series analysis, cross-spectral coherence values were used to determine whether the same temporal structure was present in the DLC and Fastrak recordings (Gottman, 1981; Warner, 1998). Zero-lag cross-correlation values were also calculated to determine the degree of sequential covariance between the DLC and Fastrak recordings (Stergiou, 2004; Winter, 2009).

We treated our data as single measurements and computed bias and limits of agreement as outlined in Bland & Altman (1986, 1999). Note that despite having multiple trials of the same task, the data cannot be considered repeated measures because the freehand motion used in this task does not follow the exact same trajectory on every trial. When calculating RMSE, we considered Fastrak data as ‘observed’ and DLC data as ‘predicted’. The cross spectral coherence and cross-correlation analysis was conducted using MATLAB 2019b. No pre-processing or data smoothing was applied to the data prior to analysis. Cross spectral coherence measures the relationship or correlation between two time-series with respect to frequency and results in a value ranging from 0 to 1, where 1 corresponds to perfect spectral coherence. Here we report the average coherence calculated at each signal’s peak frequency, with each signal’s frequency spectrum calculated using MATLABs Fast Fourier Transform (FFT) function. Cross correlation measures the covariance between two time-series signals across a specific range of temporal lags. Given that we were interested in the sequential dependence of the two signals we calculated the zero-lag correlations between the Fastrak and DLC time-series (note that the maximum cross correlation was always at a lag of 0 or 1, with the latter due to trivial delays in data synchronization). Like a standard correlation, cross-correlations values range between -1 and 1, with values close to 1 indicating a high degree of (positive) sequential covariance (i.e., signal similarity).

## Supplementary materials

The main aim of the study was to evaluate agreement between DLC and Fastrak. However, the process of obtaining 3D data from DLC involves multiple steps, each potentially introducing errors and noise. To investigate this, we ran a series of static tasks designed to reveal the influence of likely sources of noise in our DLC data: human error in labelling, network noise, camera lens distortions, and calibration inaccuracies.

### 2D static task

This task was designed to evaluate potential distortions coming from the camera lens, noise coming from different people labelling the frames, and random differences in the training process. We filmed a precisely manufactured ChArUco board (Garrido-Jurado et al. 2014) with known dimensions (Fig S1a), positioned ∼55cm away and directly facing the camera. We filmed 20 trials, 1 second each, at 120fps. All recordings were merged into a single video file for easier processing. Three different people (DK, LL, VK) independently labelled a random selection of frames. The corners of five squares on the board (four approximately in the corners, one in the center, see Fig S1b) were labelled, with 20 labelled points in total.

We ran the network training process three separate times on each set of labelled frames, obtaining nine datasets. For each dataset, we calculated the Euclidean length of each square side in pixels, based on X and Y coordinates provided by DLC. Since the ChArUco board was directly facing the camera and labelling was performed on an object with known dimensions including a known fixed side length of each individual square (10mm), we can reasonably expect a fixed true value of the square side length in pixels as well.

**Figure S1:**
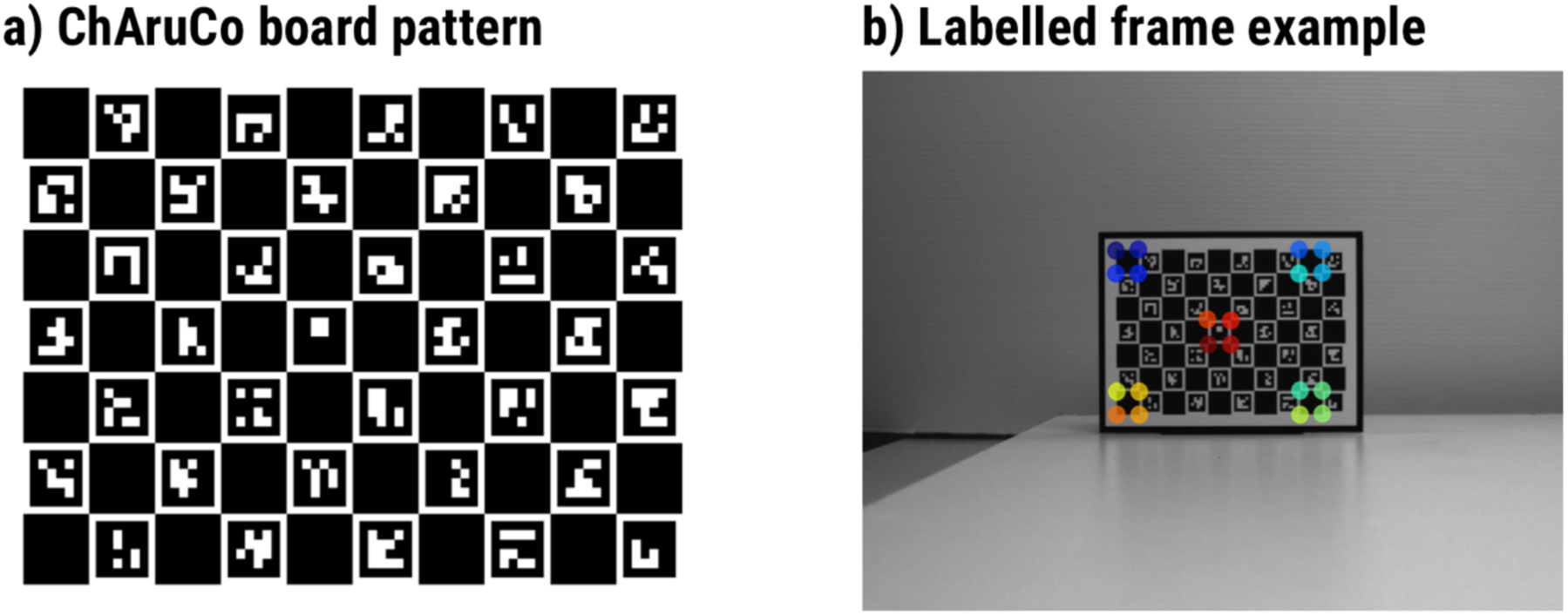
(A) Pattern printed on the ChAruCo board used for 2D and 3D static tasks. (B) An example labelled frame from 2D static task, showing the points labelled for both 2D and 3D static tasks.

### 3D static task

This task was designed to verify that the process of translating 2D data into 3D coordinates is reliable and agrees with ground truth throughout the filming volume used in the main experimental tasks (3D dynamic tasks). We filmed a stationary ChArUco board (same as above) positioned in 6 different locations in the filming space: at the front, middle, and back of the field of view, on either the left or the right half of it. The filming space and the cameras were set up identically to the 3D dynamic tasks, and we filmed 20 trials of 1 second each. We labelled the same 20 points as in the 2D static task. The calibration video was filmed during the same session as the other 3D static videos. The calibration process was the same as in the 3D dynamic task. After getting 3D coordinates for this dataset, we calculated Euclidian lengths of square sides, in millimeters.

### Results

For each static task, we calculated the mean and standard error (SE) of the square side lengths, for the relevant unit of analyses. In the 2D static task, we compared mean lengths between labelers and network training repeats. The differences between labelers were less than one pixel (see Table S1 and Figure S2a).

**Table S1:** Mean (SE) of square side lengths, in pixels, from 2D-static test.

| Network training repeat | Labeller |  |  |
| --- | --- | --- | --- |
|  | DK | LL | VK |
| 1 | 22.78 (0.003) | 23.64 (0.006) | 23.34 (0.003) |
| 2 | 22.79 (0.004) | 23.44 (0.010) | 23.56 (0.003) |
| 3 | 22.78 (0.004) | 23.62 (0.009) | 23.43 (0.003) |

The differences between network repeats were negligibly small. In the 3D static task, the worst distortion occurred at the back left of the filming space, but even there, DLC underestimates length by less than half a millimeter. DLC and the calibration process also appear to be highly reliable, as standard errors are extremely small (Table S2 and Figure S2b).

**Figure S2:**
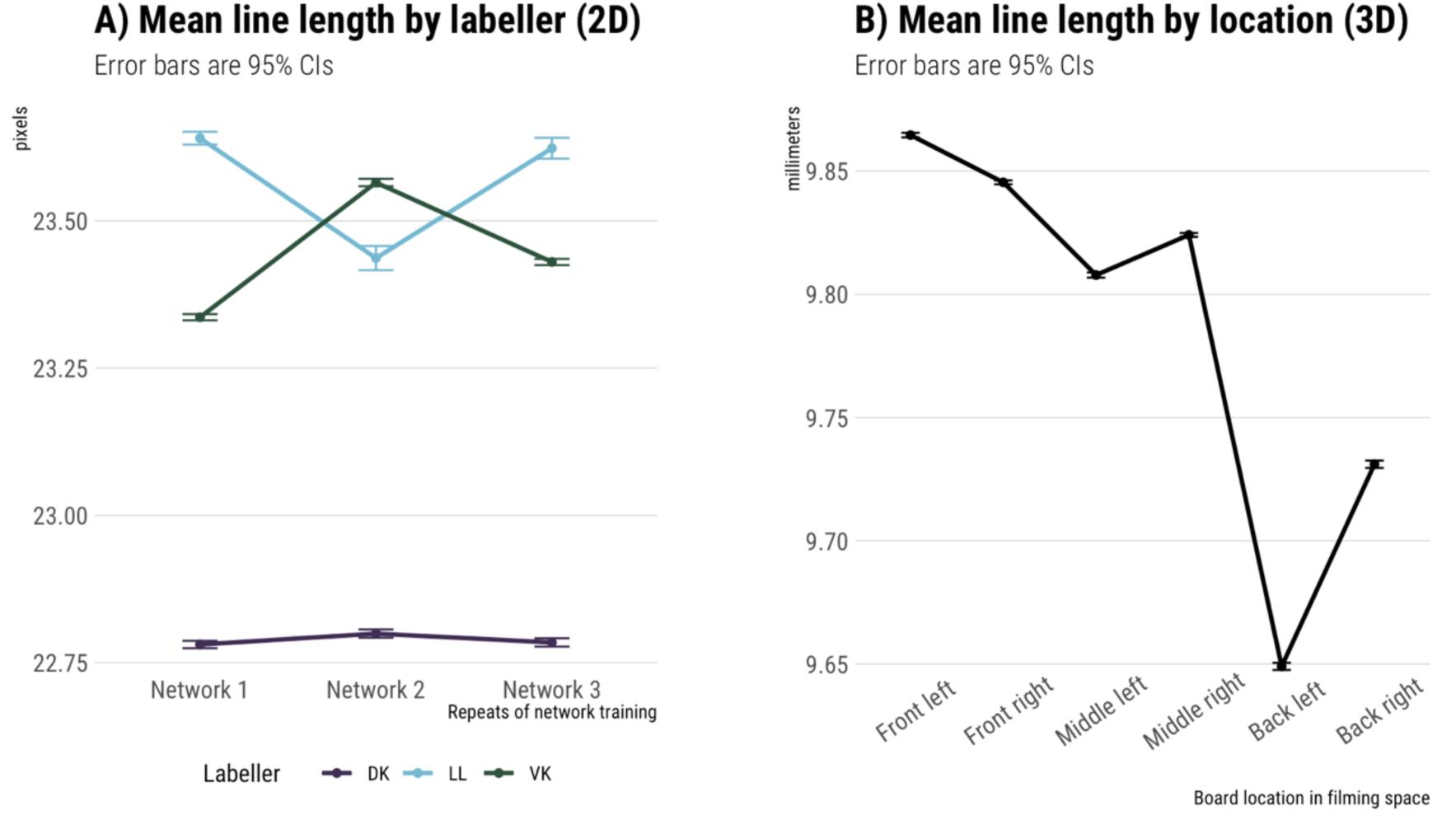
(A) Mean line length (in pixels), for each labeler and network training repeat. (B) Mean line length (in millimeters) in each of the six locations within the filming volume.

**Table 2:** Mean (SE) of square side lengths, in millimeters, at different filming locations in 3D-static test.

| Position in filming space | Left | Right |
| --- | --- | --- |
| Front | 9.86 (0.0005) | 9.85 (0.0004) |
| Middle | 9.81 (0.0005) | 9.82 (0.0004) |
| Back | 9.65 (0.0007) | 9.73 (0.0008) |

